# Quaternary structure and activity of glutamate dehydrogenase are regulated by reversible S-palmitoylation and mitochondrial acyl-protein thioesterases

**DOI:** 10.64898/2026.03.25.714181

**Authors:** Michael Salsaa, Huu Tram A Huynh, Charneal L. Dixon, Jonathan St-Germain, Haggag S. Zein, Brian Raught, Gregory D. Fairn

## Abstract

Glutamate dehydrogenase (GDH) is a key mitochondrial enzyme that catalyzes the reversible oxidative deamination of glutamate to α-ketoglutarate, thereby linking amino acid and carbohydrate metabolism. GDH forms catalytically active hexamers and is regulated by various allosteric modulators, including ADP and GTP. Here, we demonstrate that GDH undergoes auto-palmitoylation in the presence of palmitoyl-CoA, leading to a dose-dependent inhibition of enzymatic activity. Using acyl-PEG exchange assays and mass spectrometry, we show that GDH monomers are predominantly mono-palmitoylated, with modification detected at multiple cysteine residues, including Cys55, Cys115, and Cys197, among the six cysteines in the mature enzyme. Blue Native PAGE analysis revealed that palmitoylation disrupts the native hexameric assembly of mammalian GDH, which is organized as a dimer-of-trimers, promoting dissociation into dimers. Importantly, this modification is reversible, as incubation with mitochondrial acyl-protein thioesterases 1 (APT1) and, to a lesser extent, α/β hydrolase domain 10 (ABHD10) restores both the hexameric structure and enzymatic activity. The modified Cys55 residues are positioned near the trimer interface, providing a mechanism by which palmitoylation could prevent hexamer formation, whereas Cys115 and 197 may destabilize individual trimers. These findings establish *S-*palmitoylation as a novel regulatory mechanism for GDH, linking mitochondrial lipid metabolism to the reversible control of a central metabolic enzyme.

## 1 INTRODUCTION

Mitochondrial metabolism is orchestrated by a complex network of enzymes that integrate signals from multiple metabolic pathways^1, 2^. At the intersection of amino acid and carbohydrate metabolism lies glutamate dehydrogenase (GDH), an enzyme that catalyzes the reversible oxidative deamination of L-glutamate to α-ketoglutarate, producing NADH and ammonia^3^. This reaction is essential for amino acid catabolism, ammonia detoxification, and anaplerotic replenishment of tricarboxylic acid (TCA) cycle intermediates^4, 5^. GDH is ubiquitously expressed and is particularly important roles in tissues with high metabolic demands, including the liver, kidney, and brain^5^.

The quaternary structure of GDH has been extensively characterized. The enzyme functions as a homohexamer, with each subunit harboring a catalytic site^6^. The GDH hexamer consists of two trimers arranged back-to-back, with extensive inter-trimer contacts stabilizing the assembly^6, 7^. GDH activity is subject to complex allosteric regulation by metabolites including GTP, ATP and palmitoyl-CoA (inhibitors), as well as ADP and leucine (activators), allowing the enzyme to respond dynamically to the cellular energy state and metabolic flux^6, 8^. Additional layers of regulation include hormonal control and tissue-specific expression of different GDH isoforms^7, 9, 10^.

*S-*palmitoylation, the reversible attachment of palmitate to cysteine residues via thioester linkages, is an increasingly recognized post-translational modification that regulates diverse cellular processes^11^. Unlike other lipid modifications, such as myristoylation and prenylation, *S-*palmitoylation is reversible and can be dynamically regulated by cellular signals. Palmitoylation is catalyzed by a family of zinc-dependent DHHC-domain containing palmitoyltransferases (ZDHHCs)^12, 13^. At the same time, depalmitoylation is mediated by acyl-protein thioesterases, including the APT/LYPLA family and α/β-hydrolase domain-containing (ABHD) proteins^14, 15^. Recent proteomic studies have revealed that *S-*palmitoylation is far more widespread than previously appreciated, with hundreds of palmitoylated proteins identified across different cellular compartments, including mitochondria^16–18^.

Interestingly, some proteins can undergo auto-palmitoylation in the presence of palmitoyl-CoA without requiring ZDHHC enzymes, suggesting that the nucleophilicity of certain cysteine residues and their local microenvironment can facilitate direct acyl transfer^19–21^. This ZDHHC-independent mechanism may be particularly relevant in the mitochondrial matrix, where palmitoyl-CoA is generated during fatty acid β-oxidation, and ZDHHC enzymes are less abundant or absent compared to other cellular compartments^22^. Additionally, the pH of the mitochondrial matrix is typically 7.5 – 8.2^23, 24^, approaching the pKa of cysteine sulfhydryl groups, which would favour auto-palmitoylation.

GDH is known to be inhibited by palmitoyl-CoA^6^, but whether this inhibition is due to allosteric binding or to auto-palmitoylation remains unresolved^25, 26^. Given the presence of multiple cysteine residues in GDH, the comparatively high pH, and the presence of palmitoyl-CoA in mitochondria during fatty acid oxidation, we hypothesized that GDH might be subject to auto-palmitoylation and that this modification could inhibit its activity and/or quaternary structure. Here, we report that GDH undergoes auto-palmitoylation on three Cys residues, leading to hexamer dissociation and loss of enzymatic activity, and that this modification is reversible by mitochondrial-resident acyl-protein thioesterases. These findings suggest auto-palmitoylation may represent a mechanism linking lipid metabolism to the regulation of a central metabolic enzyme.

## 2 RESULTS

### 2.1 Palmitoyl-CoA inhibits glutamate dehydrogenase activity through auto-palmitoylation

We first re-examined the effect of palmitoyl-CoA on enzyme activity. Bovine liver GDH was pre-incubated with increasing concentrations of palmitoyl-CoA (4 and 10 µM) for 10 minutes, followed by measurement of enzymatic activity in the glutamate-forming (reductive amination) direction using NADH oxidation as the readout (Figure 1A). Palmitoyl-CoA treatment resulted in dose-dependent inhibition of GDH activity, with ≈35% inhibition at 4 µM and ≈85% inhibition at 10 µM palmitoyl-CoA (Figure 1B). This inhibition was not due to direct competition with substrates or cofactors, as the palmitoyl-CoA was present only during the pre-incubation period and was diluted 100-fold upon addition to the reaction mixture. This finding is also consistent with previous studies demonstrating that several long acyl-CoAs can inhibit GDH, while CoA itself and short acyl-CoAs do not^26^.

**Figure 1.**
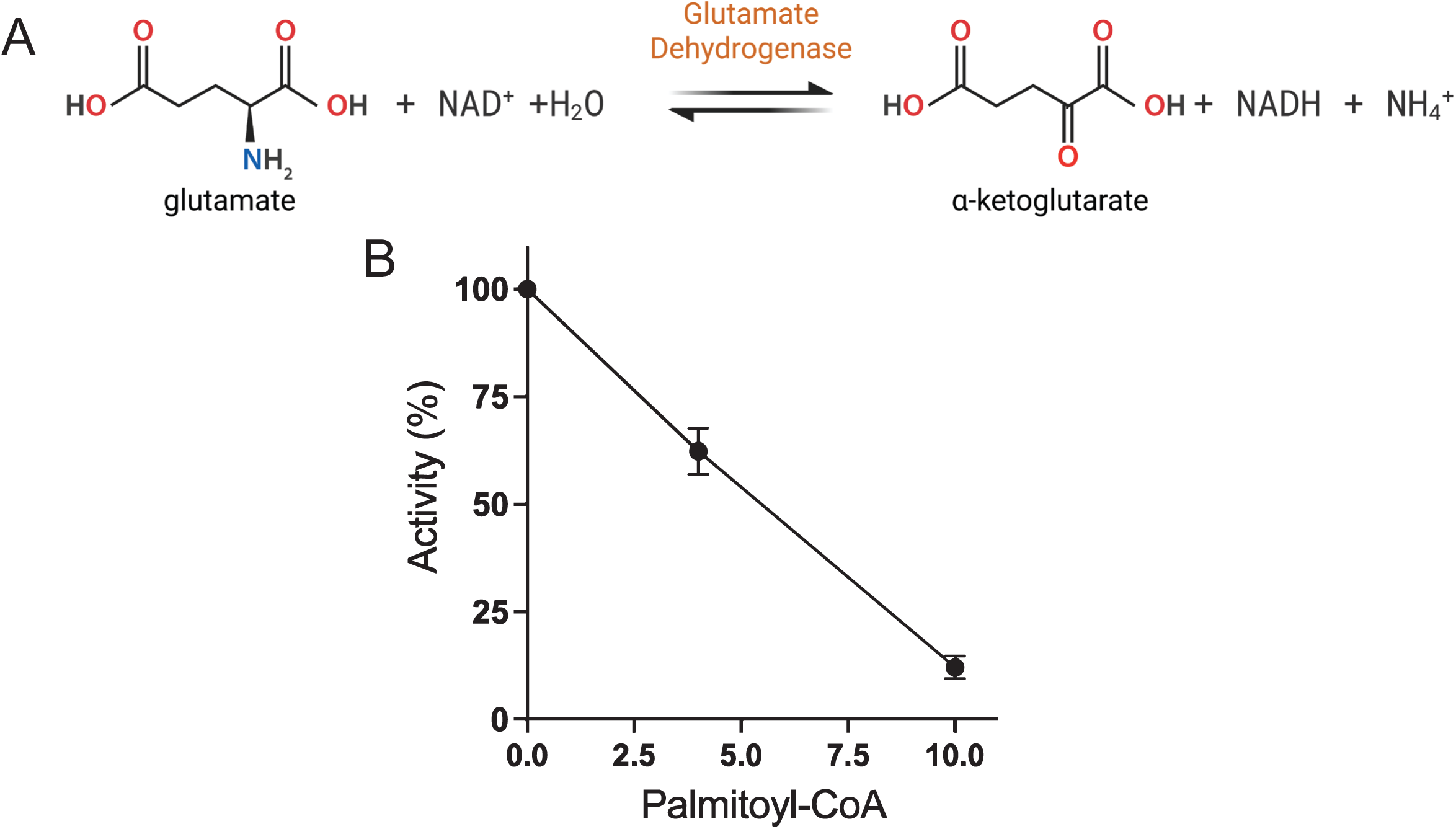
Palmitoyl-CoA inhibits glutamate dehydrogenase (GDH) activity in vitro. (A) Schematic representation of the reversible reaction catalyzed by GDH. GDH activity was measured in the glutamate-forming (reductive amination) direction, using NADH oxidation as the readout. (B) Enzymatic activity of GDH following pre-incubation with palmitoyl-CoA. GDH was treated with 0, 4, or 10 µM palmitoyl-CoA for 10 min prior to activity measurement. NADH oxidation at 340 nm was used to monitor initial rates. Data represent three independent experiments.

We suspected that in the presence of 10 μM palmitoyl-CoA at pH 8.0, deprotonated cysteine residues could act as nucleophiles and attack the carbonyl group in palmitoyl-CoA (Fig. 2A). Thus, auto-palmitoylation, rather than allosteric regulation, could explain the inhibition of GDH. To confirm that the observed inhibition resulted from covalent modification of GDH via S-acylation, we used an acyl-polyethylene glycol exchange (acyl-PEG) assay^27^. This method involves blocking free cysteines with a non-reducible alkylating agent, *S-*methyl methanethiosulfonate (MMTS), followed by selective cleavage of thioester bonds with hydroxylamine (HAM), and subsequent labeling of the newly exposed cysteines with a 5 kDa polyethylene glycol (PEG) mass tag. Palmitoylated proteins exhibit a characteristic molecular weight shift on SDS-PAGE that is dependent on hydroxylamine, which removes acyl chains from Cys residues^27^. To ensure complete auto-palmitoylation GDH was incubated with an excess of palmitoyl-CoA (1 mM). As a result, GDH showed a hydroxylamine-dependent band shift from ≍60 kDa to ≍65 kDa with the occasional appearance of a weak ≍70k Da (Figure 2B; three biological replicates run on the same gel for illustrative purposes). Quantification of the mono-PEGylated and Apo bands revealed significant hydroxylamine-dependent modification, confirming that GDH undergoes auto-palmitoylation (Figure 2C). This pattern of mono- and di-palmitoylated GDH findings suggests site-specific modifications since the mature form of bovine GDH contains six cysteine residues Cys55, Cys89, Cys115, Cys197, Cys270, and Cys319 (positions following removal of mitochondrial targeting sequence).

**Figure 2.**
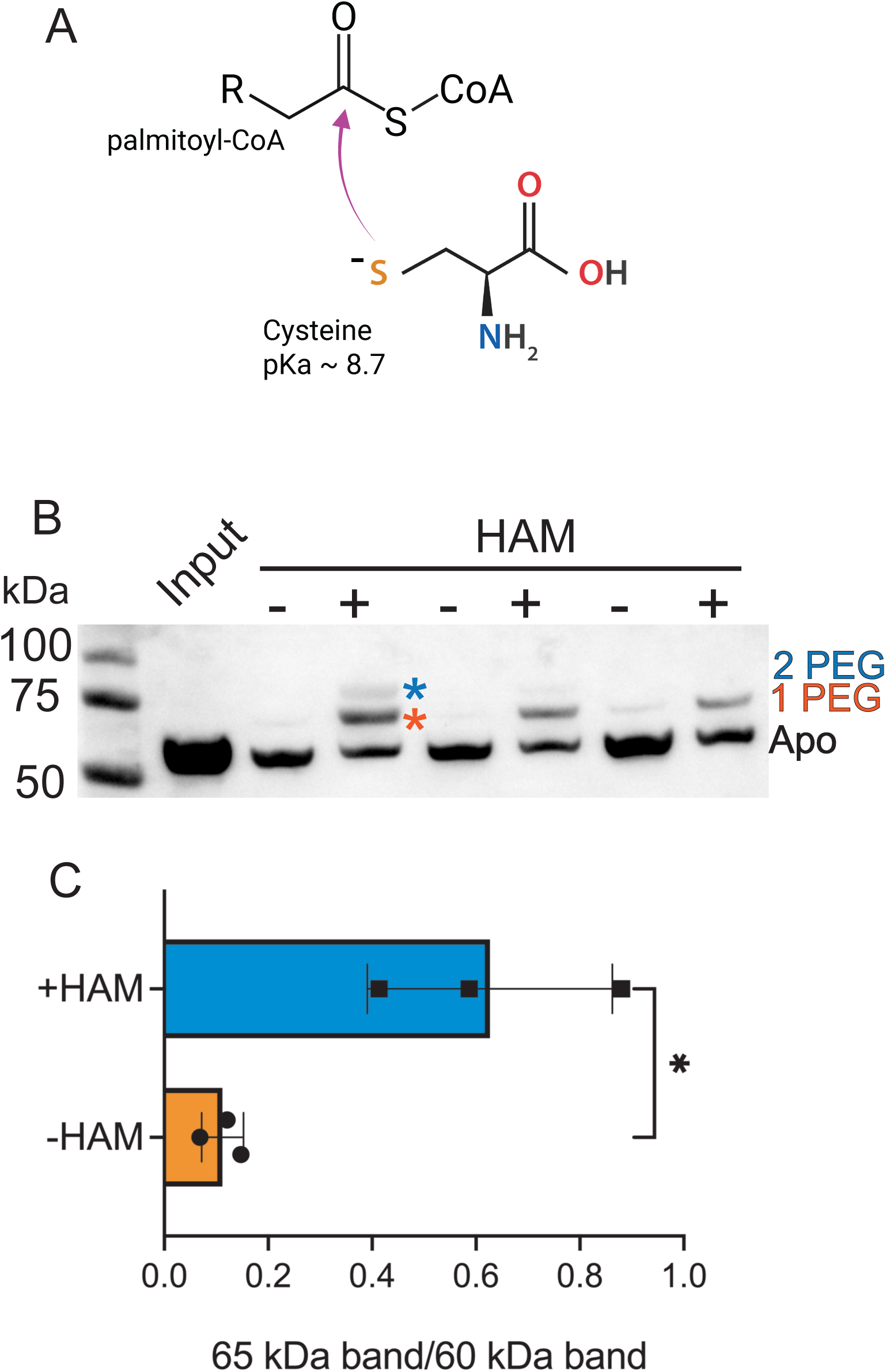
The acyl-PEG exchange assay demonstrates that GDH is predominantly mono-auto-palmitoylated. (A) Nucleophilic attack of deprotonated cysteine on palmitoyl-CoA. (B) Acyl-PEG exchange assay demonstrating palmitoylation-dependent cysteine modification. Hydroxylamine-dependent band shifts indicate the presence of thioester-linked palmitate on GDH. (C) Quantification of PEG-shifted bands from panel C was performed using AzureSpot Pro software. Data represent the means ± Std. dev. from 3 individual experiments. Statistical significance was determined by a 2-way paired t-test. Differences were considered significant for *P* ≤ .05 (*).

### 2.2 Mass spectrometry identifies multiple palmitoylation sites

To identify which cysteine residue(s) are palmitoylated, we used differential labeling and an acyl-switch strategy coupled with mass spectrometry. In this approach, palmitoylated GDH was reacted with MMTS (methylthiolation, +46 Da) to modify free cysteines. Next, the thioester bonds were cleaved with HAM and the newly exposed cysteines were alkylated with iodoacetamide. This results in carbamidomethylation (+57 Da) specifically at sites that were palmitoylated initially. LC-MS/MS analysis identified peptides containing all six cysteine residues in the mature GDH sequence (Supplemental Table 1), demonstrating good coverage of cysteine-containing regions. For reference, the fragment spectra for the relevant peptides with either IAA- or MMTS-modified peptides are included in Figure 3. Analysis of MMTS-modified and IAA-modified peptides revealed carbamidomethylation at Cys55, Cys115, and Cys197 in hydroxylamine-treated samples (Figure 4A, B). Cys55 yielded the highest spectral counts among the modified residues, though differential ionization efficiencies of distinct peptides preclude precise quantitative comparisons between sites. However, Cys89, Cys270, and Cys319 showed little to no detectable modifications under these conditions. The identification of a limited number of palmitoylation sites is consistent with our acyl-PEG exchange data, which showed primarily mono-palmitoylation with trace amounts of di-palmitoylation.

**Figure 3.**
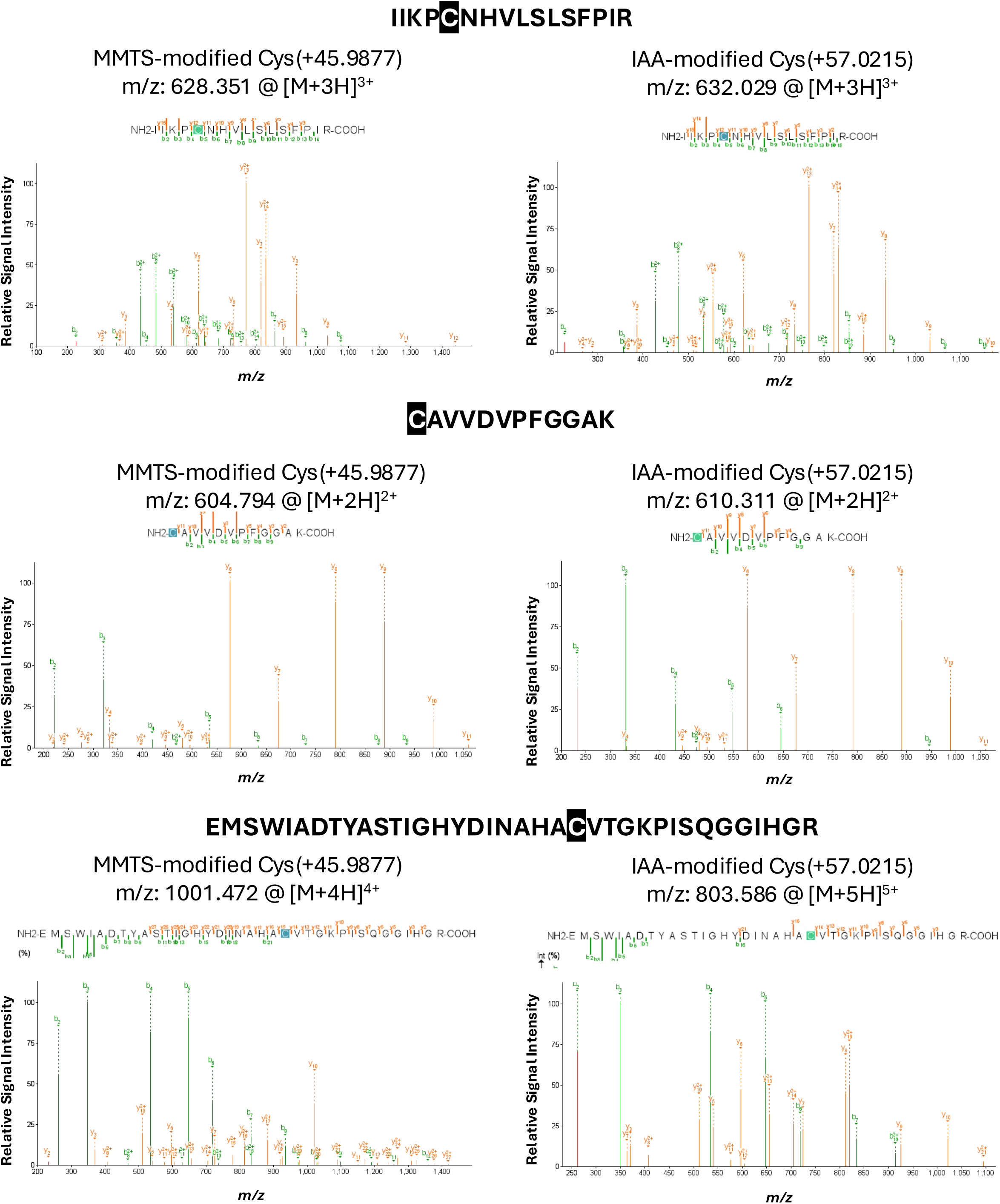
Mass spectrometry identifies palmitoylated cysteine residues in glutamate dehydrogenase using an acyl-switch strategy. Fragmentation spectra of Cys-containing bovine GDH tryptic peptides were identified as either MMTS-modified or IAA-modified.

**Figure 4.**
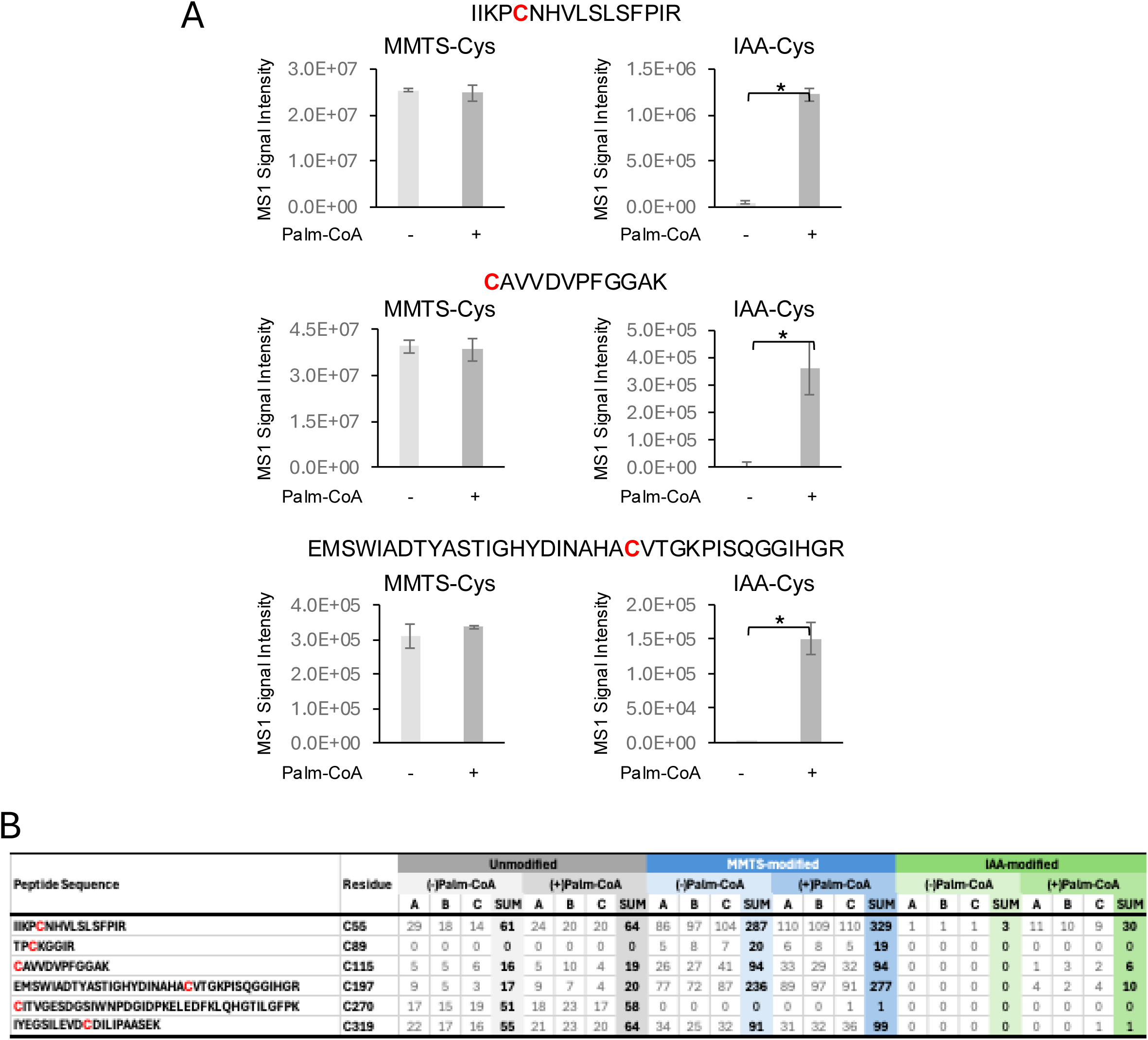
Relative proportions of glutamate dehydrogenase palmitoylated peptides. (A) MS1 signal intensities of Cys-containing glutamate dehydrogenase tryptic peptides identified as either MMTS- or IAA- modified in samples treated with or without palmitoyl-coenzyme A. (B) Peptide-spectral matching (PSM) counts of Cys-containing GDH peptides identified by LC-MS/MS. Shown are counts of unmodified Cys-containing peptides, MMTS-modified peptides (free Cys blocked with MMTS prior to hydroxylamine treatment) and iodoacetamide-modified peptides (Cys blocked with IAA following hydroxylamine treatment) in samples +/- palmitoyl-coenzyme A.

### 2.3 Structural analysis reveals palmitoylation sites are near the core at the trimer interface

To understand how palmitoylation at Cys55, Cys89 and Cys197 might affect GDH structure and function, we examined the three-dimensional structure of the bovine GDH1 hexamer using the previously published structures PBD:1NR7^28^ and PBD:3ETD bound with the inhibitor bithionol^29^. The mature human GDH hexamer forms back-to-back trimers, with extensive inter-subunit contacts stabilizing the hexameric assembly (Figure 5A). This arrangement generates three distinct types of subunit interfaces: intra-trimer contacts between adjacent subunits within each trimeric ring, intra-dimer contacts between vertically stacked subunit pairs across the two trimers, and the broader trimer-trimer interface formed by the collective back-to-back interactions.

**Figure 5.**
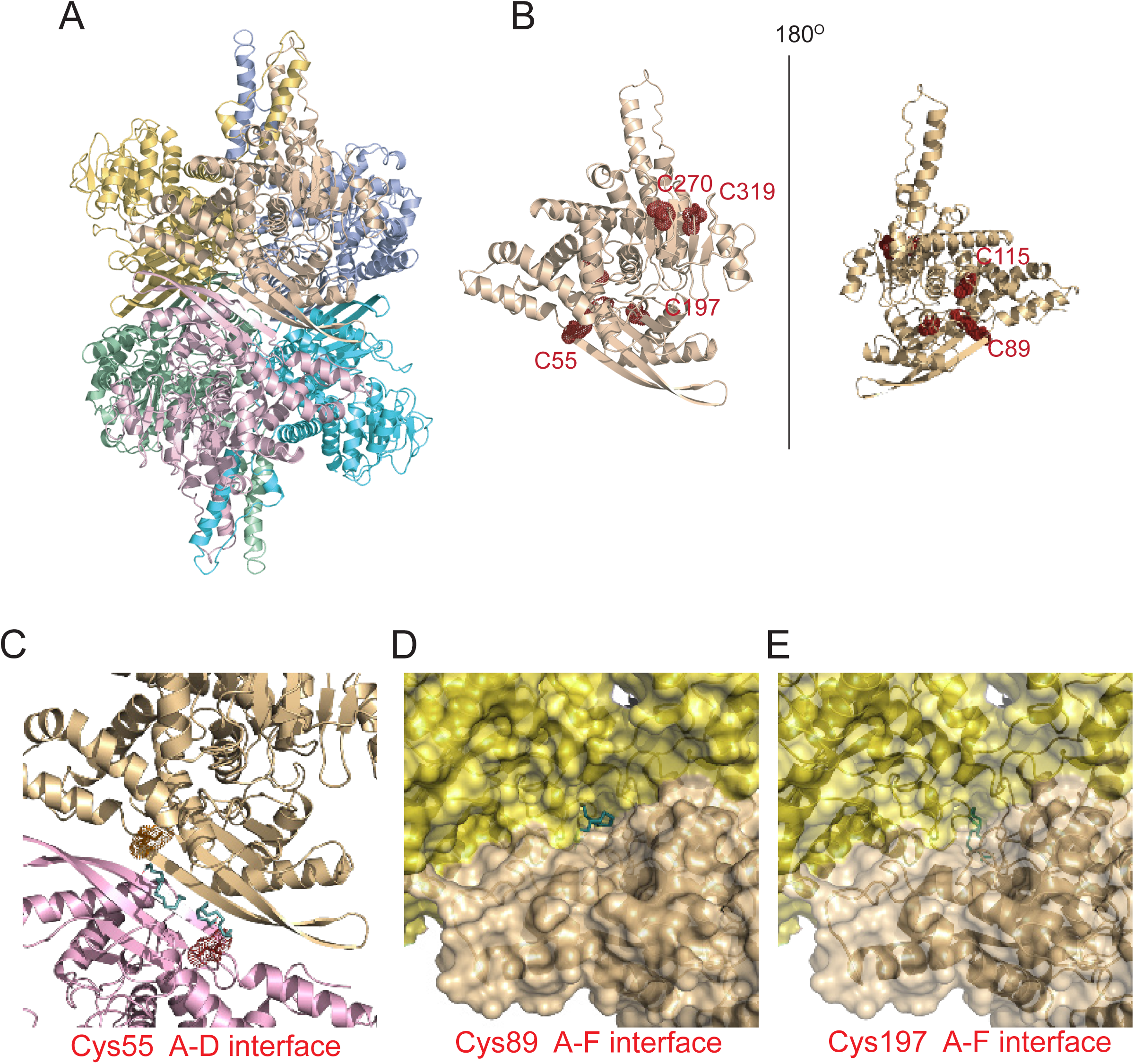
Glutamate dehydrogenase hexameric structure with bound palmitoyl groups. (A) Ribbon representations of human GDH were generated in PyMOL (version 3.1.3.1). (B) Ribbon depiction of a single GDH subunit in two orienttations highlighting the relative positions of the Cys residues. (C) shows the interface between subunits A and D with Cys55 of both being modified with palmitate. (D) shows Cys89 at the subunit A and F interface modified with palmitoyl group. (E) shows Cys197 deeper in the same pocket as Cys89 at the subunit A and F interface modified with palmitoyl group.

Mapping the positions of all six cysteine residues onto the structure revealed striking differences in their spatial locations (Figure 5B). Cys55 is located within the glutamate-binding domain (amino acids 50 – 150) at the interface where subunits from opposing trimers contact one another. In contrast, Cys274 and Cys319 are positioned on the exterior surface of the hexamer, fully exposed to solvent and distant from any subunit contact regions. Cys89, Cys115, and Cys197 occupy intermediate positions with varying degrees of surface accessibility but are not situated at critical assembly interfaces.

The location of Cys55 at the subunit A-D interface suggests a clear mechanism by which palmitoylation could disrupt hexamer stability. The trimer-trimer interface is tightly packed, with inter-subunit distances that accommodate precise complementary surfaces but leave little room for additional bulk. The attachment of a 16-carbon palmitoyl chain (∼20 Å in length) at Cys55 would introduce substantial steric bulk directly into this interface. Given the proximity of two Cys55 in the corresponding subunits, palmitoylation of both residues would introduce substantial bulk and hydrophobicity (Figure 5C). Cys89 and Cys197, the other palmitoylated residues, reside in a pocket near the interface between the A and F subunits (Figure 5D, E).

### 2.4 *S-*palmitoylation disrupts GDH hexamer formation

To directly assess the effect of palmitoylation on GDH quaternary structure, we analyzed palmitoylated GDH using Blue Native PAGE (BN-PAGE), which separates protein complexes under non-denaturing conditions while preserving native oligomeric states^30, 31^. Untreated GDH migrated predominantly as a hexamer, with an apparent molecular weight of approximately 360 kDa (Figure 6A, 0 µM lane), consistent with the enzyme’s known hexameric structure. Treatment with increasing concentrations of palmitoyl-CoA (2.5, 5, and 10 µM) resulted in a dose-dependent decrease in the hexamer population and a concomitant increase in species migrating at ≍120 kDa, consistent with dimers (Figure 6A). Occasionally, we also observed a band at 240 kDa, which may represent a dimer of dimers or a tetrameric intermediate.

**Figure 6.**
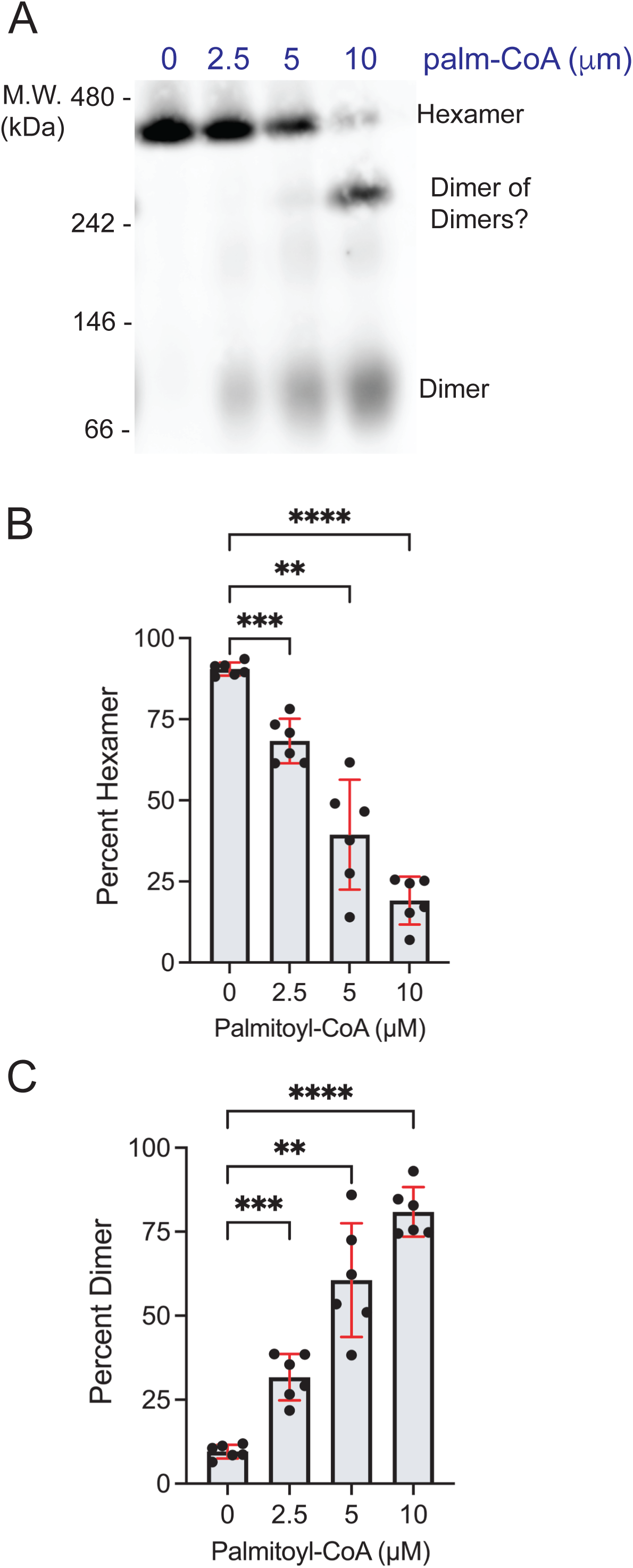
Auto-palmitoylation of glutamate dehydrogenase disrupts hexamer formation. (A) Representative Blue Native–PAGE immunoblot showing GDH oligomeric states following palmitoyl-CoA treatment. Purified GDH (10 mg/mL) was incubated with 0, 2.5, 5, or 10 µM palmitoyl-CoA for 10 min, diluted 1:10, mixed with 4× NativePAGE sample buffer, and resolved by BN-PAGE. After transfer, membranes were probed with anti-GDH antibody. (B, C) Quantification of hexamer and dimer abundances from three independent experiments. Band intensities were measured using AzureSpot Pro software. Data represent the means ± Std. error from 6 independent experiments. Statistical significance was determined by 1-way ANOVA with Tukey’s post-hoc test. Differences were considered significant for *P* ≤ .01 (**), *P* ≤ .005 (***), and *P* ≤ .001 (****).

Quantification of band intensities from three independent experiments confirmed that palmitoyl-CoA treatment caused a significant, dose-dependent decrease in the hexamer population (Figure 6B). At 10 µM palmitoyl-CoA, hexamer levels were reduced to approximately 20% of control, with a reciprocal increase in dimer formation (Figure 6C). The correlation between hexamer dissociation and loss of enzymatic activity (Figure 1B) strongly supports that the hexameric state is required for optimal GDH catalytic function and that *S-*palmitoylation-induced oligomer disruption is the primary mechanism of enzyme inhibition.

### 2.5 Acyl-protein thioesterases reverse GDH *S-*palmitoylation and restore structure and activity

*S-*palmitoylation is a reversible modification that can be removed by acyl-protein thioesterases (APTs) and α/β-hydrolase domain-containing (ABHD) enzymes^14^. APT1 is a well-characterized depalmitoylating enzyme that localizes to mitochondria and regulates mitochondrial protein depalmitoylation^32^. To test whether GDH palmitoylation is reversible, we co-incubated palmitoylated GDH with recombinant APT1 (also known as LYPLA1) or its catalytically inactive S119A mutant^32^. BN-PAGE analysis revealed that wild-type APT1 treatment restored the hexameric structure of palmitoylated GDH, while the catalytically inactive S119A mutant had no effect on oligomeric state (Figure 7A). Albumin present in the sample served as an internal loading control, confirming equal protein loading across conditions. Consistent with this, enzymatic assays showed that APT1 significantly reduced the inhibitory effect of palmitoyl-CoA on GDH activity at both 4 µM and 10 µM. In contrast, the S119A mutant did not restore activity (Figure 7B). Together, these results indicate that APT1 effectively depalmitoylates GDH, restoring both its quaternary structure and enzyme activity.

**Figure 7.**
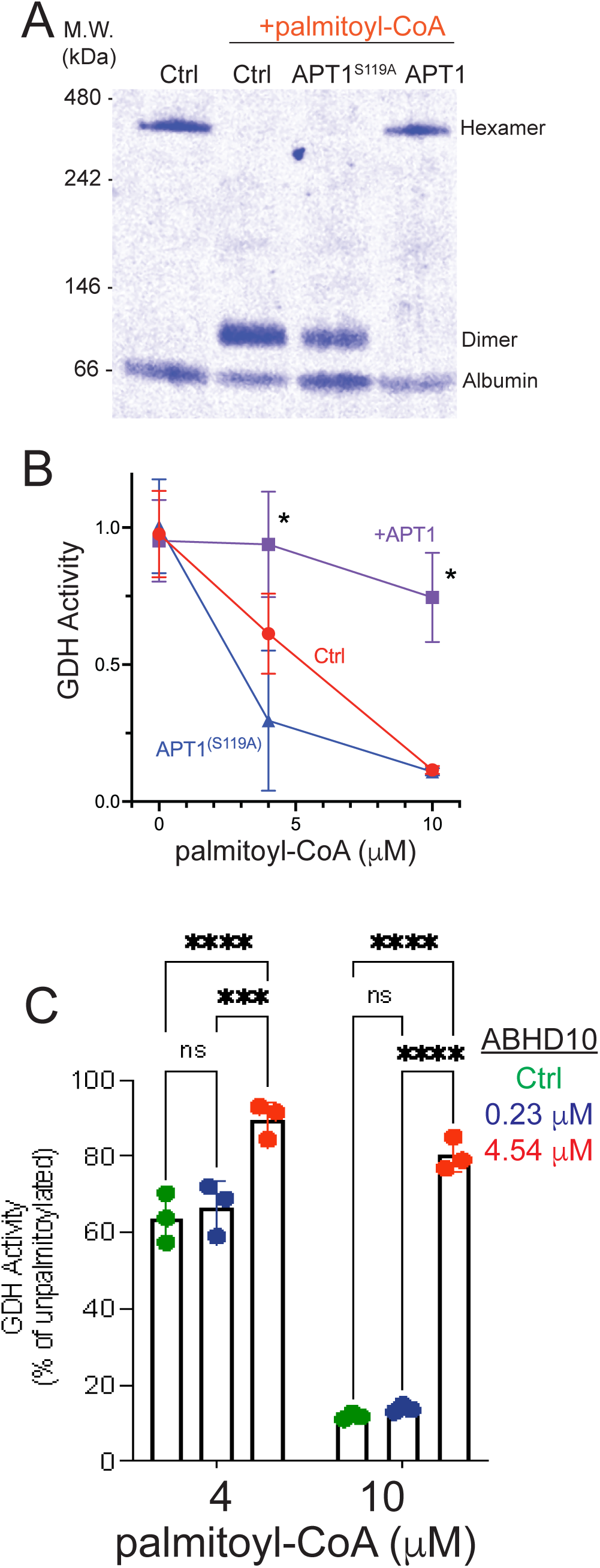
Mitochondrial acyl-protein thioesterases APT1 and ABHD10 reverse GDH auto-palmitoylation, restoring quaternary structure and enzymatic activity. (A) Blue Native–PAGE analysis of palmitoylated GDH following incubation with recombinant wild-type APT1 or the catalytically inactive S119A mutant. GDH was first auto-palmitoylated, then treated with the indicated enzymes before BN-PAGE analysis. Gels were stained with Coomassie; a representative image is shown. (B) GDH activity following palmitoylation and treatment with either APT1 or S119A. Enzyme activity was assayed in the presence of 0, 4, or 10 µM palmitoyl-CoA together with the indicated thioesterase. Activities were normalized to the untreated GDH sample (0 µM palmitoyl-CoA), which was set to 100%. Data represent three biological replicates with two technical replicates each. (C) GDH activity following palmitoylation and incubation with recombinant ABHD10. Blue circles represent ABHD10 added at the same concentration used for APT1 in panel B; red circles represent ABHD10 added at a 20-fold higher amount. Activities are shown relative to the untreated GDH control (100%). In B and C, the data represent the means ± Std. error from 3 independent experiments. Statistical significance was determined by 2-way ANOVA with Tukey’s post-hoc test. Differences were considered significant for *P* ≤ .05 (*), *P* ≤ .005 (***), and *P* ≤ .001 (****).

We next examined ABHD10, another mitochondrial depalmitoylating enzyme implicated in regulating mitochondrial protein acylation^33^. At the same concentration used for APT1, ABHD10 produced only a modest recovery of GDH activity (Figure 7C, blue circles). However, increasing ABHD10 concentration 20-fold led to substantial restoration of activity (Figure 6C, red circles), suggesting lower catalytic efficiency compared with APT1, although we can’t rule out the possibility that the recombinant ABHD10 is only partly active. Overall, these findings support a model in which GDH palmitoylation is dynamically regulated by mitochondrial acyl-protein thioesterases. The balance between palmitoylation and depalmitoylation may therefore serve as a regulatory mechanism for tuning GDH activity in response to changes in mitochondrial palmitoyl-CoA levels.

## 3 DISCUSSION

In this study, we have identified *S-*palmitoylation as a post-translational modification of glutamate dehydrogenase that regulates both its quaternary structure and catalytic activity. Our findings reveal several key insights: (1) GDH undergoes auto-palmitoylation in the presence of palmitoyl-CoA at pH 8.0, predominantly on three Cys residues; (2) palmitoylation disrupts the native hexameric structure of the enzyme, promoting dissociation into dimers; (3) the structural disruption correlates with dose-dependent inhibition of enzymatic activity; and (4) the modification is reversible by mitochondrial acyl-protein thioesterases APT1 and ABHD10. These results establish a potential link between mitochondrial lipid metabolism and the regulation of a central metabolic enzyme. In addition, our findings provide mechanistic insight into the long-standing observation that long-chain acyl-CoAs inhibit GDH *in vitro*.

The mechanism by which *S-*palmitoylation of Cys55, disrupts hexamer formation can be rationalized by previously determined structures. Cys55 lies within the glutamate-binding domain (residues ∼50-150), which forms the core of the hexamer and mediates the critical trimer-trimer interface through extensive antiparallel β-sheet contacts between vertically stacked subunit pairs^29^. During normal catalysis, this core region undergoes a “breathing” motion in which stacked dimer pairs move 2.5-4% closer together as the catalytic cleft closes, then separate as it opens^29^. This dynamic process is essential for catalytic turnover—the stacked dimers must be able to move relative to each other for efficient substrate binding and product release. The attachment of a 16-carbon palmitoyl chain at Cys55 would introduce ≈21 Å of hydrophobic bulk directly into this interface^34^. Unlike non-covalent inhibitors such as bithionol and GW5074, which bind to the nearby α-helix (residues 138-155) and inhibit GDH by restricting the breathing motion without causing disassembly^29^, the covalent nature of palmitoylation creates a permanent steric barrier. The results would predict that the palmitoyl chain cannot be accommodated by the normal dynamic range of the interface and thus forces complete separation of the trimer-trimer contacts. This interface—formed by the extensive antiparallel β-sheet between vertically stacked subunits—remains essentially unchanged during catalysis (∼0.1% distance variation) and represents the tightest and most stable association within the hexamer. Cys55 is positioned to disrupt the lateral contacts between these stacked dimers, not the vertical contacts within them. Consequently, palmitoylation at Cys55 would specifically weaken the trimer-trimer interface while leaving the intra-dimer β-sheet contacts intact, causing the hexamer in trimers. However, our BN-PAGE results detect primarily dimers, not trimers. Thus, it is conceivable that palmitoylation of Cys89 or Cys197 disrupts the interactions within the trimer.

Our finding that hexamer dissociation correlates with loss of enzymatic activity is consistent with previous studies demonstrating that the hexameric state is required for optimal GDH function^6, 35^. Importantly, the reversibility of this process by APT1 and ABHD10 demonstrates that palmitoylation-mediated regulation is dynamic and potentially responsive to changes in cellular metabolic state. The physiological relevance of GDH palmitoylation remains to be fully established, but several metabolically coherent scenarios can be envisioned. Under high-glucose conditions, mitochondrial acetyl-CoA is derived primarily from glycolysis, whereas during fasting or low-glucose states, fatty acids become the dominant energy source and long-chain acyl-CoAs such as palmitoyl-CoA are transported into the mitochondrial matrix for β-oxidation. Elevated mitochondrial acyl-CoA availability under these conditions may promote GDH auto-palmitoylation and suppression of amino acid oxidation. Such regulation would be metabolically advantageous, as fatty acid β-oxidation and amino acid catabolism compete for shared limiting cofactors, including NAD⁺, FAD, CoASH, and electron transport chain capacity. Because GDH consumes NAD⁺ and generates NADH during glutamate oxidation, excessive GDH activity could impair β-oxidation by elevating the NADH/NAD⁺ ratio. In this context, palmitoylation-mediated inhibition of GDH may preserve NAD⁺ availability and prioritize fatty acid oxidation during lipid-dependent metabolic states.

The identification of APT1 and ABHD10 as enzymes capable of depalmitoylating GDH is also noteworthy. Both enzymes have been localized to mitochondria and are emerging regulators of mitochondrial protein acylation. APT1 showed robust activity toward palmitoylated GDH, whereas ABHD10 required higher concentrations to achieve similar effects, suggesting differences in substrate specificity or catalytic efficiency. However, we can’t rule out the possibility that the recombinant bacterially expressed ABHD10 may not be fully active. Regardless, the presence of multiple depalmitoylating enzymes with overlapping but distinct specificities may provide redundancy and fine-tuning of GDH palmitoylation dynamics in response to different metabolic signals.

Our findings expand our understanding of GDH regulation beyond classical allosteric mechanisms and support a paradigm for metabolic enzyme control through reversible lipid modification. More broadly, our results highlight the importance of considering S-Palmitylation/Acylation as a regulatory mechanism for enzymes, particularly in mitochondria, where acyl-CoA species are abundant and metabolic pathways must be precisely coordinated. Recent studies have demonstrated that S-acylation of the active site of patatin-like phospholipase domain-containing protein 2 (PNPLA2) is essential for its catalytic activity, representing a direct role in enzyme function^36^. Our findings here add another dimension to this emerging paradigm by demonstrating that S-acylation can also function as an inhibitory modification by disrupting quaternary structure. These examples highlight that S-acylation should be considered as a versatile regulatory mechanism capable of modulating protein function through multiple distinct mechanisms: membrane targeting, catalytic enhancement, and allosteric inhibition via structural remodeling. The growing recognition of S-acylation as a dynamic and functionally diverse post-translational modification suggests that many metabolic enzymes may be subject to similar lipid-based regulation.

## 4 CONCLUSION

We have demonstrated that glutamate dehydrogenase undergoes reversible *S-*palmitoylation at Cys55, Cys89, and Cys197, thereby disrupting hexamer formation and inhibiting enzymatic activity. This palmitoylation is reversed by mitochondrial acyl-protein thioesterases APT1 and ABHD10, suggesting a dynamic regulatory mechanism linking mitochondrial lipid metabolism to amino acid catabolism. These findings reveal a previously unrecognized layer of GDH regulation and suggest that palmitoylation may serve as a broader mechanism for coordinating metabolic flux in response to changes in cellular energy status and nutrient availability.

## 5 MATERIALS AND METHODS

### 5.1 Materials

Bovine liver glutamate dehydrogenase was obtained from Roche (Cat. 10197734001) as a powder and resuspended in 50 mM triethanolamine (TEA), pH 8.0, at 10 mg/mL. Palmitoyl-CoA (Sigma, Cat. P9716) was resuspended in methanol at 10 mM. All other reagents were of analytical grade and purchased from Sigma-Aldrich unless otherwise specified.

### 5.2 Plasmids and constructs

Expression plasmids encoding human APT1 (LYPLA1), catalytically inactive APT1 (S119A), and ABHD10 were generously provided by Bryan Dickinson (Department of Chemistry, University of Chicago) and are based on the pET-30a vector^32, 33^.

### 5.3 *In vitro* GDH activity assay

GDH activity was measured by monitoring NADH oxidation at 340 nm using a SpectraMax i3X plate reader (Molecular Devices, USA) at 25°C. Reactions were carried out in a final volume of 200 µL and contained 50 mM triethanolamine buffer (pH 8.0), 100 mM ammonium acetate, 0.1 mM NADH, 2.6 mM EDTA, 8 mM α-ketoglutarate, and GDH at 10 µg/mL. The decrease in absorbance at 340 nm was monitored for 10 minutes, and initial velocities were calculated from the linear portion of each reaction trace. One unit of enzyme activity was defined as the amount of protein required to oxidize 1 µmol of NADH per minute at 25°C.

For palmitoylation experiments, purified GDH was pre-incubated with palmitoyl-CoA at the indicated concentrations for 10 minutes at room temperature, while control samples were treated with an equal volume of methanol. Following this pre-incubation, the enzyme was added to the reaction mixture described above, and activity was recorded immediately. All experiments were performed in three independent replicates, each with two technical replicates.

### 5.4 Blue Native PAGE and immunoblotting

The oligomeric state of GDH was analyzed using Blue Native PAGE (BN-PAGE). Samples were prepared in 4× NativePAGE™ Sample Buffer (Invitrogen, Cat. BN20032) and loaded onto 4-16% NativePAGE™ Bis-Tris gels (Invitrogen, Cat. BN2112BX10). Electrophoresis was performed using NativePAGE™ Running Buffers (Invitrogen, Cat. BN2001 and BN2002). A NativeMark™ unstained protein standard was included as a molecular weight reference. Gels were run at 150 V for the first hour and increased to 250 V for an additional hour at room temperature.

Following electrophoresis, proteins were either stained with Coomassie overnight or transferred to PVDF membranes using NuPAGE™ Transfer Buffer (Invitrogen, Cat. NP0006) at 4°C for 2 hours by wet transfer. After transfer, membranes were incubated in 8% acetic acid for 15 minutes, rinsed with deionized water, air-dried completely, and briefly soaked in methanol before additional washing in water. Membranes were then blocked in 5% nonfat milk in TBST for 1 hour and probed with anti-GLUD1 primary antibody (Proteintech, Cat. 14299-1-AP) overnight, followed by an IRDye-700-conjugated goat anti-rabbit secondary antibody (Cat. AC2128). Fluorescence imaging was performed using an Azure 600 imaging system (Azure Biosystems).

### 5.5 Protein expression and purification

Recombinant proteins were expressed in *Escherichia coli* Rosetta™ 2 (DE3) pLysS cells (Novagen). Starter cultures were prepared by inoculating 30 mL LB medium containing 50 µg/mL kanamycin and incubating overnight at 37°C with shaking at 240 rpm. The overnight cultures were then used to inoculate 500 mL LB supplemented with 50 µg/mL kanamycin, which was grown at 37°C and 240 rpm to an OD of 0.6-0.8. At mid-log phase, cultures were chilled on ice and induced with 100 µM IPTG. Protein expression proceeded at 10°C with shaking at 240 rpm for 20 hours. Cells were harvested by centrifugation at 4,000 x g for 15 minutes and stored at -20°C until purification.

Cell pellets were resuspended in 10 mL B-PER™ bacterial extraction reagent (Thermo Fisher Scientific) supplemented with DNase I (Roche), incubated for 15 minutes at room temperature, and clarified by centrifugation at 15,000 × g for 15 minutes at 4°C. The supernatant was incubated with cobalt affinity resin (Co²D-NTA, Thermo Fisher Scientific, Cat. 25229) at 4°C overnight with gentle rotation. Beads were washed five times with wash buffer (50 mM HEPES, pH 8.0, 150 mM NaCl), and bound protein was eluted with 300 mM imidazole in 50 mM HEPES, pH 8.0.

### 5.6 Acyl-PEG exchange assay

Acyl-PEG exchange was performed to assess the S-acylation state of GDH. Commercial GDH (Sigma, Cat. G2501) was diluted to 0.5 mg/mL in 50 mM triethanolamine (TEA) buffer, pH 8.0. Palmitoylation was initiated by adding palmitoyl-CoA to a final concentration of 1 mM and incubating the mixture at room temperature for 10 minutes. For thiol blocking, 100 µL of enzyme solution from each replicate was mixed with 100 µL blocking buffer (100 mM HEPES, pH 7.5, 1 mM EDTA, 2.5% SDS) containing 1% MMTS (Sigma, Cat. 208795). Samples were incubated at 40°C with shaking at 1,500 rpm for 1 hour.

Blocked samples were then split into two equal fractions and precipitated using the methanol-chloroform method. Pellets were washed with methanol, air-dried, and resuspended in 30 µL binding buffer (100 mM HEPES, pH 7.5, 1 mM EDTA, 1% SDS). Thioester cleavage was performed by adding 30 µL of 1 M neutral hydroxylamine (NHDOH) to one fraction, while the corresponding control received 30 µL of water. Samples were incubated at room temperature for 2 hours with shaking at 1,500 rpm. Proteins were again precipitated, washed, and resuspended in 60 µL binding buffer containing 2 mM methoxypolyethylene glycol maleimide (mPEG-maleimide; Sigma, Cat. 63187). PEGylation proceeded for 2 hours at room temperature with shaking at 1,500 rpm. Following the final precipitation and methanol wash, pellets were resuspended in 4× loading buffer and analyzed by SDS-PAGE.

### 5.7 Acyl-switch assay for mass spectrometry

GDH powder was reconstituted in 50 mM triethanolamine (TEA) buffer, pH 8.0, at a concentration of 10 µg/µL. For palmitoylation, 30 µL of GDH was incubated with 60 µL of 1 mM palmitoyl-CoA for 10 minutes at room temperature. Control samples were treated with an equal volume of methanol. Free cysteine residues were then blocked by adding each sample to 300 µL of blocking buffer containing 15 µL MMTS, followed by incubation at 40°C for 1 hour at 1,500 rpm.

Proteins were precipitated with 1.1 mL ice-cold acetone and washed three times with 70% acetone. After air-drying, pellets were resuspended in 200 µL binding buffer and briefly sonicated. Thioester cleavage was performed by adding 100 µL of 1 M neutral hydroxylamine and 5 mM iodoacetamide, and samples were incubated in the dark at room temperature with gentle rotation for 2 hours. Samples were then submitted for LC-MS/MS analysis.

Samples were lyophilized and reconstituted in 50 mM NH_4_HCO_3_ prior to the addition of 1 µg of trypsin. Following an overnight incubation at 37□, samples were acidifed with HCOOH (1% final concentration), desalted on C18 tips, and lyophilized. Peptides were reconstituted in HCOOH (0.1% final concentration) and 500ng of sample were loaded onto an Evotip Pure™ column (Evosep, Odense). Liquid chromatography was performed with the Evosep One (Evosep, Odense Denmark) pump with an SPD30 method using an Evosep Performance C18 column (EV1137, 15 cm x 150 µm ID, 1.5 µm; Evosep, Odense). The TIMS-TOF HT (Bruker, Bremen) mass spectrometer was operated in PASEF-DDA positive ion mode (MS scan range 100-1700 m/z). Ion mobility range was 1/K0 = 1.6 to 0.6 Vs cm-2 using equal ion accumulation and ramp time in the dual TIMS analyzer of 100 ms each. Collision energy was lowered stepwise as a function of increasing ion mobility, starting from 20 eV for 1/K0 = 0.6 Vs cm-2 and 59 eV for 1/K0 = 1.6 Vs cm-2. The ion mobility dimension was calibrated linearly using three ions from the Agilent ESI LC/MS tuning mix (m/z, 1/K0: 622.0289, 0.9848 Vs cm-2; 922.0097, 1.1895 Vs cm-2; and 1221.9906, 1.3820 Vs cm-2). Data files were analyzed on the Fragpipe platform (v22.0) using MSFragger^37, 38^. The UniprotKB bovine reference proteome (Bos taurus; UP000009136; 22,666 entries) was searched with trypsin set as the digestion enzyme (2 missed cleavage allowed), variable modifications used were +57.0215 and +45.9877 at Cys residues to account for iodoacetamide and MMTS modifications, respectively, in addition to acetylation (protein N-term), oxidation (M), and deamidation (NQ). Search engine results were filtered with Philosopher^39^, Percolator^40^ and Protein Prophet^41^.

### 5.8 Molecular Docking

Molecular docking simulations were performed using AutoDock Vina (version 1.5.6)^42^ to investigate the binding orientation of palmitic acid within the active site of mature glutamate dehydrogenase (GDH). The crystal structure of bovine GDH (PDB ID: 3ETD) was retrieved from the Protein Data Bank. Prior to docking, all crystallographic water molecules and co-crystallized ligands (GTP, NADPH, glutamate, and bithionol) were removed. Polar hydrogen atoms were added, and Kollman charges were assigned using AutoDockTools (ADT). Palmitic acid (PLM) was obtained from the Protein Data Bank entry and prepared by defining rotatable bonds and saving the structure in PDBQT format. Docking simulations were performed by centering the search space on each of the six cysteine residues (Cys55, Cys89, Cys115, Cys197, Cys270, and Cys319) of monomer A within the GDH hexamer. In addition, docking was performed at Cys55 of monomer D to evaluate the orientation of palmitate at the monomer A–D interface. A grid box with dimensions of 20 Å × 20 Å × 20 Å was defined for each docking run, centered to encompass the target cysteine residue. Docking calculations were executed with an exhaustiveness value of 8, generating 9 poses per run. The resulting docking poses were ranked based on predicted binding affinity (kcal/mol), and the top-ranked conformation (lowest binding energy) for each site was selected for further interaction analysis. Observed binding energies of top-ranked poses ranged from −5.8 to −3.3 kcal/mol. Protein–ligand complexes and hydrogen bonding interactions were visualized using PyMOL (version 3.1.3.1) from Shrödinger.

### 5.9 Statistical analysis

All data are presented as mean ± standard deviation from at least three independent experiments. Statistical significance was assessed using 1- or 2-way ANOVA followed by Tukey’s post-hoc test for multiple comparisons or Student’s *t*-test for pairwise comparisons. Statistical significance was defined as *P* ≤ 0.05 (*), *P* ≤ 0.01 (**), *P* ≤ 0.005 (***), *P* ≤ 0.0001 (****). All analyses were performed using GraphPad Prism 9.0.

## AUTHOR CONTRIBUTIONS

M.S. designed and performed experiments, analyzed data, and wrote the manuscript. A.H. and C.D. performed experiments and analyzed data. J.S.-G. and B.R. performed mass spectrometry analysis and wrote the manuscript. G.D.F. conceived the study, designed experiments, analyzed data, and wrote the manuscript. All authors reviewed and approved the final manuscript.

## ACKNOWLEDGEMENTS

We thank Bryan Dickinson (University of Chicago) for generously providing expression plasmids for APT1, ABHD10, and related constructs. This work was funded by the Canadian Institutes of Health Research – Institute of Cancer Research (CR3 – 190819 to G.D.F.) and Grant 1058003 from the Cancer Research Society (to G.D.F.). G.D.F. is also supported by a Tier 1 Canada Research Chair in Multiomics of Lipids and Innate Immunity. A.H. received funding from the Beatrice Hunter Cancer Research Institute through the Canadian Cancer Society’s Carol Ann Cole Comfort Heart for Breast Cancer Research. Work in the Raught lab was supported by the Philip S. Orsino Chair in Leukemia Research, a joint Hospital-University Named Chair between the University of Toronto, The Princess Margaret Cancer Centre Director, and the Princess Margaret Cancer Foundation.

## REFERENCES

1. Spinelli, J.B. & Haigis, M.C. The multifaceted contributions of mitochondria to cellular metabolism. Nat Cell Biol 20, 745–754 (2018).

2. Martinez-Reyes, I. & Chandel, N.S. Mitochondrial TCA cycle metabolites control physiology and disease. Nat Commun 11, 102 (2020).

3. Hudson, R.C. & Daniel, R.M. L-glutamate dehydrogenases: distribution, properties and mechanism. Comp Biochem Physiol B 106, 767–792 (1993).

4. Plaitakis, A. & Zaganas, I. Regulation of human glutamate dehydrogenases: implications for glutamate, ammonia and energy metabolism in brain. J Neurosci Res 66, 899–908 (2001).

5. Spanaki, C. & Plaitakis, A. The role of glutamate dehydrogenase in mammalian ammonia metabolism. Neurotox Res 21, 117–127 (2012).

6. Li, M., Li, C., Allen, A., Stanley, C.A. & Smith, T.J. The structure and allosteric regulation of mammalian glutamate dehydrogenase. Arch Biochem Biophys 519, 69–80 (2012).

7. Peterson, P.E. & Smith, T.J. The structure of bovine glutamate dehydrogenase provides insights into the mechanism of allostery. Structure 7, 769–782 (1999).

8. Smith, T.J. & Stanley, C.A. Untangling the glutamate dehydrogenase allosteric nightmare. Trends Biochem Sci 33, 557–564 (2008).

9. Plaitakis, A., Metaxari, M. & Shashidharan, P. Nerve tissue-specific (GLUD2) and housekeeping (GLUD1) human glutamate dehydrogenases are regulated by distinct allosteric mechanisms: implications for biologic function. J Neurochem 75, 1862–1869 (2000).

10. Mastorodemos, V., Zaganas, I., Spanaki, C., Bessa, M. & Plaitakis, A. Molecular basis of human glutamate dehydrogenase regulation under changing energy demands. J Neurosci Res 79, 65–73 (2005).

11. Chamberlain, L.H. & Shipston, M.J. The physiology of protein S-acylation. Physiol Rev 95, 341–376 (2015).

12. Mitchell, D.A., Vasudevan, A., Linder, M.E. & Deschenes, R.J. Protein palmitoylation by a family of DHHC protein S-acyltransferases. J Lipid Res 47, 1118–1127 (2006).

13. Greaves, J. & Chamberlain, L.H. DHHC palmitoyl transferases: substrate interactions and (patho)physiology. Trends Biochem Sci 36, 245–253 (2011).

14. Won, S.J., Cheung See Kit, M. & Martin, B.R. Protein depalmitoylases. Crit Rev Biochem Mol Biol 53, 83–98 (2018).

15. Lin, D.T. & Conibear, E. ABHD17 proteins are novel protein depalmitoylases that regulate N-Ras palmitate turnover and subcellular localization. Elife 4, e11306 (2015).

16. Sanders, S.S. et al. Curation of the Mammalian Palmitoylome Indicates a Pivotal Role for Palmitoylation in Diseases and Disorders of the Nervous System and Cancers. PLoS Comput Biol 11, e1004405 (2015).

17. Kostiuk, M.A. et al. Identification of palmitoylated mitochondrial proteins using a bio-orthogonal azido-palmitate analogue. FASEB J 22, 721–732 (2008).

18. Dennis, K. & Heather, L.C. Post-translational palmitoylation of metabolic proteins. Front Physiol 14, 1122895 (2023).

19. Duncan, J.A. & Gilman, A.G. Autoacylation of G protein alpha subunits. J Biol Chem 271, 23594–23600 (1996).

20. Guan, X. & Fierke, C.A. Understanding Protein Palmitoylation: Biological Significance and Enzymology. Sci China Chem 54, 1888–1897 (2011).

21. Dietrich, L.E. & Ungermann, C. On the mechanism of protein palmitoylation. EMBO Rep 5, 1053–1057 (2004).

22. Ohno, Y., Kihara, A., Sano, T. & Igarashi, Y. Intracellular localization and tissue-specific distribution of human and yeast DHHC cysteine-rich domain-containing proteins. Biochim Biophys Acta 1761, 474–483 (2006).

23. Llopis, J., McCaffery, J.M., Miyawaki, A., Farquhar, M.G. & Tsien, R.Y. Measurement of cytosolic, mitochondrial, and Golgi pH in single living cells with green fluorescent proteins. Proc Natl Acad Sci U S A 95, 6803–6808 (1998).

24. Santo-Domingo, J. & Demaurex, N. Perspectives on: SGP symposium on mitochondrial physiology and medicine: the renaissance of mitochondrial pH. J Gen Physiol 139, 415–423 (2012).

25. Son, H.J. et al. Roles of cysteine residues in the inhibition of human glutamate dehydrogenase by palmitoyl-CoA. BMB Rep 45, 707–712 (2012).

26. Kawaguchi, A. & Bloch, K. Inhibition of glutamate dehydrogenase and malate dehydrogenases by palmitoyl coenzyme A. J Biol Chem 251, 1406–1412 (1976).

27. Percher, A. et al. Mass-tag labeling reveals site-specific and endogenous levels of protein S-fatty acylation. Proc Natl Acad Sci U S A 113, 4302–4307 (2016).

28. Banerjee, S., Schmidt, T., Fang, J., Stanley, C.A. & Smith, T.J. Structural studies on ADP activation of mammalian glutamate dehydrogenase and the evolution of regulation. Biochemistry 42, 3446–3456 (2003).

29. Li, M., Smith, C.J., Walker, M.T. & Smith, T.J. Novel inhibitors complexed with glutamate dehydrogenase: allosteric regulation by control of protein dynamics. J Biol Chem 284, 22988–23000 (2009).

30. Schagger, H. & von Jagow, G. Blue native electrophoresis for isolation of membrane protein complexes in enzymatically active form. Anal Biochem 199, 223–231 (1991).

31. Schagger, H., Cramer, W.A. & von Jagow, G. Analysis of molecular masses and oligomeric states of protein complexes by blue native electrophoresis and isolation of membrane protein complexes by two-dimensional native electrophoresis. Anal Biochem 217, 220–230 (1994).

32. Kathayat, R.S. et al. Active and dynamic mitochondrial S-depalmitoylation revealed by targeted fluorescent probes. Nat Commun 9, 334 (2018).

33. Cao, Y. et al. ABHD10 is an S-depalmitoylase affecting redox homeostasis through peroxiredoxin-5. Nat Chem Biol 15, 1232–1240 (2019).

34. Tanford, C. The hydrophobic effect and the organization of living matter. Science 200, 1012–1018 (1978).

35. Bell, E.T. & Bell, J.E. Catalytic activity of bovine glutamate dehydrogenase requires a hexamer structure. Biochem J 217, 327–330 (1984).

36. Zheng, Y. et al. S-acylation of ATGL is required for lipid droplet homoeostasis in hepatocytes. Nat Metab 6, 1549–1565 (2024).

37. Kong, A.T., Leprevost, F.V., Avtonomov, D.M., Mellacheruvu, D. & Nesvizhskii, A.I. MSFragger: ultrafast and comprehensive peptide identification in mass spectrometry-based proteomics. Nat Methods 14, 513–520 (2017).

38. Teo, G.C., Polasky, D.A., Yu, F. & Nesvizhskii, A.I. Fast Deisotoping Algorithm and Its Implementation in the MSFragger Search Engine. J Proteome Res 20, 498–505 (2021).

39. da Veiga Leprevost, F., et al. Philosopher: a versatile toolkit for shotgun proteomics data analysis. Nat Methods 17, 869–870 (2020).

40. Kall, L., Canterbury, J.D., Weston, J., Noble, W.S. & MacCoss, M.J. Semi-supervised learning for peptide identification from shotgun proteomics datasets. Nat Methods 4, 923–925 (2007).

41. Nesvizhskii, A.I., Keller, A., Kolker, E. & Aebersold, R. A statistical model for identifying proteins by tandem mass spectrometry. Anal Chem 75, 4646–4658 (2003).

42. Trott, O. & Olson, A.J. AutoDock Vina: improving the speed and accuracy of docking with a new scoring function, efficient optimization, and multithreading. J Comput Chem 31, 455–461 (2010).

